# Long-chain Polyunsaturated Fatty Acids Mitigate *In Vitro* Skeletal Muscle Wasting Induced by Colorectal Carcinoma Cells via a 15-LOX-dependent Pathway

**DOI:** 10.64898/2026.05.22.726995

**Authors:** Xinyue Lu, Krishna Rao Maddipati, James F. Markworth

## Abstract

Up to 50% of adults with colorectal cancer (CRC) are at risk of progressive involuntary loss of skeletal muscle mass and function known as cachexia. Available options to prevent and treat cachexia in cancer survivors are currently limited. Long-chain polyunsaturated fatty acids (LC-PUFAs) and their bioactive metabolites, termed specialized pro-resolving lipid mediators (SPMs), promote the resolution of inflammation and support muscle growth and repair. However, prior studies of cachexia have mainly focused on fish oil supplements, and it is not fully understood how different individual omega-3 (n-3) and omega-6 (n-6) LC-PUFAs mediate CRC-induced muscle wasting. In addition, the crosstalk between cancer cells, the host immune system, and skeletal muscle cells in response to LC-PUFA treatments remains unclear. This study aimed to examine the effects of n-3 and n-6 LC-PUFAs on CRC-induced muscle wasting and the underlying cellular and molecular mechanisms involved. Using murine C2C12 skeletal muscle cells and CT26 colorectal carcinoma cells, we investigated the impacts of LC-PUFAs including arachidonic acid (ARA, 20:4n-6), eicosapentaenoic acid (EPA, 20:5n-3), docosapentaenoic acid (DPA, 22:5n-3), and docosahexaenoic acid (DHA, 22:6n-3) on CT26-induced muscle cell wasting in the presence or absence of lipoxygenase (LOX) inhibitors such as NDGA or BLX-3887. We also examined the lipidomic profile of C2C12-CT26 co-cultures in response to individual LC-PUFA treatments. Our results suggest that LC-PUFAs including ARA, EPA, DHA, and DPA each individually protect against CRC-induced muscle cell wasting *in vitro*, and these protective effects are dependent on 15-LOX activity. Furthermore, we found that C2C12-CT26 co-culture produced mature SPMs in response to individual PUFA treatments. Taken together, this study suggests that individual n-3 and n-6 LC-PUFAs can mitigate CRC-associated cachexia primarily by producing 15-LOX-derived bioactive lipid mediators.

## INTRODUCTION

Colorectal cancer (CRC) is the third most diagnosed cancer and the second deadliest malignancy worldwide (1). Up to 50% of colorectal cancer patients are at risk of developing cancer cachexia, a complex multifactorial syndrome characterized by progressive functional impairment resulting from loss of skeletal muscle mass and strength (2, 3). The loss of skeletal muscle mass negatively impacts the quality of life, survival rate, and tolerance of anti-cancer treatments in CRC patients (4–6). Despite active research on anti-cachectic strategies, therapeutic treatments to prevent and treat CRC cachexia are currently limited (4). As a result, there is an urgent need to develop anti-cancer therapies that both inhibit tumor growth and simultaneously preserve muscle mass to improve clinical outcomes in cancer survivors.

CRC-induced muscle wasting could be driven by multiple mechanisms such as elevated inflammation, an imbalance between catabolic and anabolic signaling, and mitochondrial dysfunction (7). Among these, chronic elevation of circulating pro-inflammatory cytokines, such as tumor necrosis factor alpha (TNFα) and interleukin- 6 (IL-6) is one of the most observed clinical features of cancer cachexia (8). Anti-inflammatory treatments such as non-steroidal anti-inflammatory drugs (NSAIDs) and selected cytokine antagonists have long been proposed as potential therapies for cancer cachexia (9, 10). Nevertheless, anti-inflammatory drugs such as corticosteroids are catabolic and immunosuppressive and thus may potentially exert negative effects on skeletal muscles in the long run (11). In addition, the complex interaction between tumor cells, skeletal muscle, and the host immune system in cachexia syndrome could limit the effectiveness of a single anti-inflammatory therapy (12).

Omega-3 (n-3) long-chain polyunsaturated fatty acids (LC-PUFAs) are natural dietary lipids with well-known anti-inflammatory properties (13). Previous research has shown that cancer-induced loss of skeletal muscle was associated with a lower circulating level of n-3 PUFAs (14, 15). Additionally, dietary supplementation with fish oils rich in n-3 LC-PUFAs can reduce circulating inflammatory markers such as IL-6 in human cancer patients (16, 17). Preclinical studies have also shown that fish oil supplementation can protect against cancer-related muscle wasting in tumor-bearing rodent models (18). Similarly, randomized, double-blinded, and placebo-controlled human trials in cancer patients have reported promising effects of dietary fish oil supplementation on maintaining body weight (19–21). Nevertheless, some other clinical trials suggested that fish oil supplementation may not be able to limit cancer-induced muscle wasting (22–24). Notably, evidence specific to CRC-related cachexia has been largely limited.

Most prior cancer cachexia research studies have evaluated fish oil supplements that contain a complex mixture of n-3 PUFAs including eicosapentaenoic acid (EPA, 20:5n-3) and docosahexaenoic acid (DHA, 22:6n- 3), amongst many others, with considerable variability in dosage and purity of PUFAs (19, 23). Differences in lipid composition, PUFA molecular structure, and dosage could potentially contribute to the inconsistent results observed across studies. Several studies have reported that the administration of pure EPA exhibits anti-cachectic effects in tumor bearing mice by inhibiting lipolysis, stimulating protein synthesis, and repressing protein degradation (25–27). While some earlier studies reported no benefits of DHA or alpha linolenic acid (ALA, 18:2n- 3) supplementation on attenuating cancer cachexia (28–30), pure DHA has more recently been reported to protect against chemotherapy-induced body weight loss in tumor-bearing rats (31). Other bioactive lipids such as the intermediary n-3 LC-PUFA docosapentaenoic acid (DPA, 22:5n-3) and the major n-6 PUFA arachidonic acid (ARA, 20:4n-6) have also been shown to potential benefit muscle health in a non-cancer environment. For example, we previously showed that supplementation of cultured muscle cells with various individual LC-PUFAs including ARA, EPA, DPA, and DHA could each individually promote C2C12 muscle cell hypertrophy (32). Additionally, ARA-derived metabolites including prostaglandin F_2α_ (PGF_2α_) can stimulate protein synthesis and promote muscle hypertrophy (33). Consistently, many studies have shown that blockade of prostaglandin biosynthesis by pharmacological inhibitory of the cyclooxygenase (COX) pathway could impair myogenesis (34). However, to date, no prior studies have compared the potential impacts of various individual n-3 and n-6 PUFAs on CRC cachexia.

One potential mechanism that may explain the anti-cachectic effects of LC-PUFAs is the downstream production of specialized pro-resolving lipid mediators (SPMs). SPMs are bioactive lipid mediators which can be derived from both n-3 and n-6 LC-PUFAs via the action of three major biosynthetic enzymes including cyclooxygenase (COX), lipoxygenase (LOX), and epoxygenase (CYP) (35). SPMs have recently been shown to repress cancer-induced inflammation and promote anti-tumor immunity through diverse mechanisms (36). For instance, resolvin D1 (RvD1) and resolvin D2 (RvD2) which are derived from n-3 DHA via the 15-LOX/5-LOX pathway could repress the secretion of tumor-induced proinflammatory cytokines and stimulate macrophage phagocytosis (37). Resolvin E1 (RvE1) derived from n-3 EPA via the coordinated action of the CYP/5-LOX pathway could reduce reduced tumor-associated neutrophils and macrophages (38). Notably, lipoxin A_4_ (LXA_4_) derived from the n-6 ARA by the 15-LOX/5-LOX pathway and/or 5-LOX/12-LOX pathway could promote anti-tumor immunity by inhibiting regulatory B cells in animal models of colorectal cancer (39). In addition, LXA_4_ treatment at a later cancer stage could increase peripheral neutrophils while suppressing lymphocytes (40). Emerging evidence indicates that SPMs modulate skeletal muscle inflammation and regeneration following acute injury (41). Moreover, growing evidence suggests that SPMs including LXA_4_, RvD1, RvD2, PD1, and RvE1 could suppress muscle inflammation and alleviate muscle wasting in health conditions such as Duchenne muscular dystrophy (42–45). Nevertheless, the potential role of SPMs in CRC-associated muscle wasting remains to be determined.

In the current study, we examined the potential effects of supplementation with various individual n-3 and n-6 LC-PUFAs on CRC-induced muscle wasting. Moreover, we sought to determine the role of different lipid mediator biosynthetic pathways in mediating the effects of LC-PUFAs on CRC cachexia. Using *in vitro* models, we examined the direct influence of n-3 and n-6 LC-PUFAs on C2C12 myotube wasting induced by tumor-secreted factors. Intriguingly, we show that individual LC-PUFAs including ARA, EPA, DPA, and DHA each alleviated CRC-induced C2C12 myotube atrophy. Furthermore, the benefits of each of these LC-PUFAs are entirely dependent on the activity of 15-LOX, a pivotal enzyme catalyzing SPM biosynthesis. To characterize the potential capacity of tumor-muscle cell crosstalk in producing SPMs, conditioned media samples derived from a cancer-muscle co-cultures receiving LC-PUFA supplementation were analyzed by LC-MS/MS-based metabolipidomic profiling. Cachectic tumor-muscle cell co-cultures receiving LC-PUFA supplementation produces many lipid metabolites, including bioactive SPMs. Moreover, direct administration of mature SPMs produced by ARA, EPA, DPA, and DHA exert differential protective effects on CRC-induced muscle wasting. In short, our results could contribute to a better understanding of how the different types of dietary LC-PUFAs may influence cancer cachexia as well as their potential mechanisms. Ultimately, this could guide personalized dietary and nutritional supplementation guideline in cancer survivors.

## METHODS

### Cell culture

The murine skeletal muscle cell line C2C12 (ATCC, CRL-1772) and colorectal carcinoma cell line CT26 (ATCC, CRL-2638) were cultured separately in growth media (GM) consisting of Dulbecco’s modified Eagle medium (DMEM, Gibco, 11995-065) containing 10% fetal bovine serum (FBS, Corning, 35-010-CV) and antibiotics (penicillin 100 U/mL, streptomycin 100 μg/mL, Gibco, 15140-122) at 37 °C in cell incubator with 5% CO_2_. To induce myotube formation confluent C2C12 myoblasts were cultured in differentiation media (DM) consisting of DMEM containing 2% horse serum (HS, Gibco 26050088) and antibiotics for 72 h prior to experimental treatments.

### LC-PUFA and SPM Treatment

Individual LC-PUFAs including ARA (20:4n-6) (Cayman Chemical, 90010), EPA (20:5n-3) (Cayman Chemical, 90110), DPA (22:5n-3) (Cayman Chemical, 90165), and DHA (22:6n-3) (Cayman Chemical, 90310) were aliquoted and stored at – 80 °C under nitrogen gas until use. Single use vials of LC-PUFAs were thawed on ice and prepared in C2C12 differentiation media. In all the experiments involving PUFA treatments, the final concentration of ARA, EPA, DPA, or DHA was 25 µM, and 0.05% ethanol in differentiation media was used as the corresponding vehicle control. Individual mature SPMs including LXA_4_ (Cayman Chemical, 90410), RvE1 (Cayman Chemical, 10007848), RvD1 (Cayman Chemical, 10012554), RvD5 (Cayman Chemical, 10007280), protectin D1 (PD1) (Cayman Chemical, 10010390), maresin 1 (MaR1) (Cayman Chemical, 10878), and n-3 DPA-derived resolvin D2 (RvD2_n-3 DPA_) (Cayman Chemical, 34482) were stored at – 80 °C under nitrogen gas until use. SPMs were prepared in C2C12 differentiation media and administered at a dose of 100 nM, with 0.28% ethanol in differentiation media used as the appropriate vehicle control.

### TNFα-induced Muscle Wasting

Subconfluent C2C12 cells were seeded into a 12-well plate in GM at a cell density of 2.5 × 10^4^/cm^2^ and allowed to proliferate and crowd in growth media for 3 days. Then, GM was replaced by DM to induce C2C12 differentiation for 72 hours. To mimic the influence of cancer-derived pro-inflammatory cytokines recombinant mouse TNFα (Cayman Chemical, 32069) was added to mature C2C212 myotubes at day 3 post-differentiation at a dose of 100 ng/mL. After the media change, individual PUFAs or SPMs were spiked into the wells as described above for a further 72 hours before fixation and immunocytochemistry.

### CT26 Conditioned media

To obtain colorectal carcinoma cell conditioned media (CM), CT26 cells were seeded at 2.67 × 10^4^/cm^2^ in a T75 flask in serum-free DMEM containing antibiotics. After 48 hours, the culture media from CT26 cells was collected and centrifuged at 500 × g to remove cell debris and the supernatant was stored at -20 °C until use. On the day of treatment, CT26 media was thawed once and mixed with DMEM containing 4% HS at a 1:1 dilution to reach a final of 50% CT26 media with 2% HS (CT26 CM). To induce muscle wasting, C2C12 myoblasts were differentiated in DM for 72 hours as described above, and mature C2C12 myotubes received fresh CT26 CM. Then, individual PUFAs or SPMs were spiked into the C2C12 myotubes as described above for another 72 hours before fixation and immunocytochemistry.

### Muscle-Cancer Co-culture

C2C12 myoblasts were seeded into a 24-well co-culture base plate (Corning, 353504) in growth media at a cell density of 2.5 × 10^4^/cm^2^ and allowed to proliferate and crowd for 72 hours. Then, growth media was replaced by DM to induce myogenic differentiation. On the same day, CT26 cells were seeded in the upper compartments of co-culture plate inserts (Corning, 353095) at a cell density of 1.67 × 10^5^/cm^2^. In parallel, 1.67 × 10^5^/cm^2^ of C2C12 cells were plated in the inserts as a non-cancer cell control for the co-culture experiment. 72 hours after C2C12 differentiation started, the upper co-culture inserts were moved to the lower compartments of the base plates. Fresh DM was added to replace culture media in both the base plates and the co-culture inserts after the compartments were moved together. Individual LC-PUFAs including ARA, EPA, DPA, and DHA were spiked into both the lower compartments and the inserts at 25 µM as described above. After 72 hours, conditioned culture media samples were collected from the lower compartments for LC-MS/MS analysis and C2C12 myotubes were fixed and stained for visualization as described below.

### LOX Inhibitors

To examine the role of LOX-derived metabolites in mediating the effects of LC-PUFAs on CT26- induced muscle atrophy, commercially available pan-LOX inhibitor nordihydroguaiaretic acid (NDGA) (Cayman Chemicals, 70300) or the 15-LOX specific inhibitor BLX3887 (Cayman Chemicals, 27391) were prepared in CT26 CM to reach a final concentration of 50 µM and 10 µM, respectively. Confluent C2C12 myoblasts were differentiated for 72 hours, after which the differentiation media was replaced by CT26 CM with or without NDGA or BLX-2887. Then, individual PUFAs including ARA, EPA, DHA, and DPA were spiked into the wells to reach a final concentration of 25 µM. Following 72 hours of incubation in the presence of experimental treatments, C2C12 myotubes were fixed in 4% paraformaldehyde (PFA) in preparation for immunocytochemistry analysis.

### Immunocytochemistry and Image Analysis

To visualize cell morphology after exposure to cancer factors, C2C12 myotubes were fixed in 4% paraformaldehyde (PFA, Electron Microscopy Sciences, 15710) for 30 minutes at 4°C and then permeabilized with 0.1% Triton X-100 for 30 minutes at room temperature. C2C12 cells were blocked in 1% bovine serum albumin (BSA, Sigma-Aldrich A3294-50G) for 1 hour at room temperature and incubated with primary antibodies against sarcomeric myosin (MF20c, DSHB, 1:100) and myogenin (F5Dc, DSHB, 1:100) prepared in blocking buffer at 4°C overnight. The next day, C2C12 cells were washed in PBS and then incubated with secondary antibodies including Goat Anti-Mouse IgG1 Alexa Fluor 568 (Invitrogen A-21124, Thermo Fisher Scientific, 1:500) and Goat Anti-Mouse IgG2b Alexa Fluor 647 (Invitrogen, Thermo Fisher Scientific A-21242, 1:500). DAPI (Invitrogen, Thermo Fisher Scientific D21490, 2 μg/mL) was used to counterstain the cell nuclei. C2C12 myotubes were then visualized with the fluorescent microscope (Echo Revolution) operating in inverted configuration. Nine images were automatically captured using a 10 × Plan Fluorite objective from the same predetermined positions in each culture well to avoid investigator bias. To assess myotube diameters, the ten largest myosin-positive cells were chosen from each image and the distance of their widest uniform point perpendicular to the length of the cell manually was measured using the straight-line selection tool in Image J/FIJI. Each branch from branching myotubes was measured as a single myotube, and the region where the branches converge was excluded from analysis.

### RNA extraction and RT-qPCR

For CT26 CM experiments, confluent C2C12 myoblasts were induced to differentiate for 48 h, and C2C12 cells were pretreated with 25 μM dose of individual PUFAs including ARA, EPA, DPA, and DHA for another 24 hours. Then, DM was replaced by CT26 CM, and a 25 μM dose of individual PUFAs including ARA, EPA, DPA, and DHA were spiked into CM for 3 hours. For SPM experiments, confluent C2C12 myoblasts were induced to differentiate for 72 h, and then myotubes received fresh DM containing 100 ng/mL of TNFα. 30 minutes after, C2C12 myotubes were then treated with 100 nM of individual SPMs including RvD1, RvD5, RvE1, RvD2_n-3 DPA_, LXA_4_, MaR1, and PD1 for 3 hours. RNA was extracted using TRIzol reagent and chloroform/isopropanol isolation. The RNA yield was determined using a NanoDrop 1000 UV/Vis Spectrophotometer (NanoDrop Technologies, E112352). Any contaminating genomic DNA was removed by Ambion™ DNase I (Invitrogen, Thermo Fisher Scientific, AM2222). RNA (2 μg) was reverse-transcribed to cDNA using High-Capacity RNA-to-cDNA Kit (Thermo Fisher Scientific 4387406). RT-qPCR was performed in 10 μL reactions of PowerUp SYBR™ Green Master Mix (Thermo Fisher Scientific, A25742) with 1 μM forward and 1 μM reverse primer (**Table 1**) on a 384-well QuantStudio™ 5 Real-Time PCR Instrument (Thermo Fisher Scientific, A28135) with samples loaded in duplicates. Relative mRNA expression fold change was calculated using the 2^−ΔΔCT^ method, with Beta-2-microglobulin (*B2m*) used as the endogenous control.

**Table 1.**
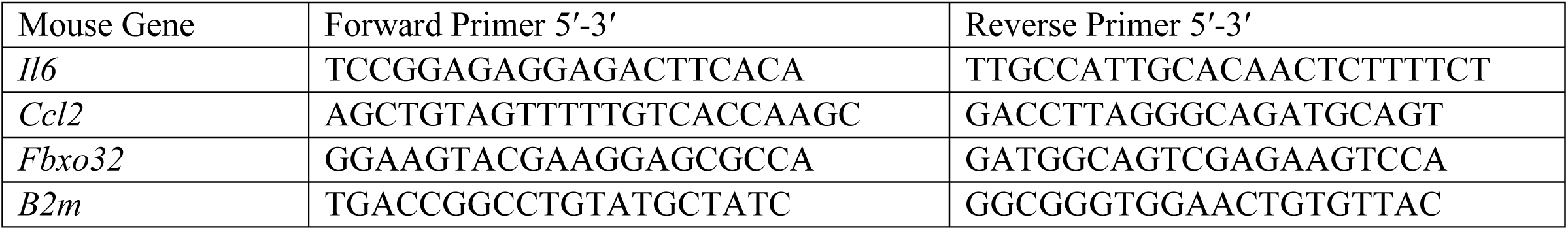
Primer Sequences for Genes used in RT-qPCR.

### LC-MS/MS-based metabolipidomic profiling of conditioned media

Conditioned cell culture media samples (500 µL) were collected from the lower compartments of the muscle-cancer co-culture model as described above and centrifuged at 500 × g for 5 min at 4 °C. Media supernatants were then collected and stored at −80 °C until Liquid Chromatography-Tandem Mass Spectrometry (LC-MS/MS) analysis.

Prior to solid phase extraction (SPE), conditioned media samples (450 μL) were spiked with 1 ng each of 15(S)- HETE-d8, 14(15)-EpETrE-d11, Resolvin D2-d5, Leukotriene B4-d4, and Prostaglandin E1-d4 as internal standards (in 150 μL methanol) for recovery and quantitation and mixed thoroughly. The samples were then extracted for polyunsaturated fatty acid metabolites using C18 extraction columns as previously described (46) Briefly, the internal standard spiked samples were applied to conditioned C18 cartridges, washed with 15% methanol in water followed by hexane, and then dried under vacuum. The cartridges were eluted with 2 volumes of 0.5 mL methanol with 0.1% formic acid. The eluate was dried under a gentle stream of nitrogen. The residue was redissolved in 50 μL methanol–25 mM aqueous ammonium acetate (1:1) and subjected to LC-MS/MS analysis.

HPLC was performed using a ExionLC-40 UPHLC system (Sciex) with Luna C18 (3 μm, 2.1 × 150 mm) column. The mobile phase consisted of a gradient between A: methanol–water-acetonitrile (10:85:5 v/v), and B: methanol– water-acetonitrile (90:5:5 v/v), both containing 0.1% ammonium acetate. The gradient program with respect to the composition of B was as follows: 0–1 min, 50%; 1–8 min, 50%–80%; 8–15 min, 80%–95%; and 15–17 min, 95%. The flow rate was 0.2 mL/min. The HPLC eluate is directly introduced to ESI source of QTRAP7500 mass analyzer (SCIEX) in the negative ion mode with following conditions: Curtain gas and GS1: 40 psi, and GS2: 70 psi, Temperature: 500 °C, Ion Spray Voltage: -2500 V, Collision gas: l2 psi, Declustering Potential: -60 V, and Entrance Potential: -7 V. The eluate is monitored by Multiple Reaction Monitoring (MRM) method to detect unique molecular ion – daughter ion combinations for each of the 125 transitions (to monitor a total of 156 lipid mediators). The MRM is scheduled to monitor each transition for 120 s around the established retention time for each lipid mediator. Optimized Collisional Energies (18 – 35 eV) and Collision Cell Exit Potentials (7 – 10 V) are used for each MRM transition. Mass spectra for each detected lipid mediator were recorded using the Enhanced Product Ion (EPI) feature to verify the identity of the detected peak in addition to MRM transition and retention time match with the standard. The data are collected and MRM transition chromatograms quantitated using Sciex OS 3.4 software. Integrated peaks with signal/noise ratio >3 were used for quantification. The internal standard signals in each chromatogram are used for normalization for recovery as well as relative quantitation of each analyte. All lipid mediators quantified were positively identified by comparing HPLC retention times with authentic standards and specific parent–daughter ion combinations as well as mass spectra obtained from information dependent acquisition (IDA) method. A full list of analytes with expected retention times, MRM transitions, and matching MRM-Internal standing mapping used for quantification is shown in Table S1A.

### Analysis for lipidomics

Analytes were classified as Not Detected (ND) in each experimental group if their peak signal-to-noise (S/N) ratios were <3 in ≥50% of samples per group. For analytes with S/N ratios ≥3 in ≥50 of samples per group, remaining missing values were replaced with half the minimal positive value in the original data set. A heatmap of the 50 most affected lipid mediators was generated in MetaboAnalyst 6.0 using the Euclidean distance measure and the Ward clustering algorithm following autoscaling of features without data transformation or normalization. Pairwise volcano plot analysis including fold changes (FC), raw p-values, and Benjamini-Hochberg (BH) false discovery rate (FDR) adjusted p-values were generated in MetaboAnalyst 6.0 following log transformation of raw data. Targeted parametric statistical analysis was performed on a predetermined subset of metabolites of interest.

### Statistical analysis

Data is shown as the mean ± SEM with dot plots displaying the raw data of three independent culture wells (considered to be biological replicates). Statistical analysis was performed in GraphPad Prism 10. Differences between treatment groups were detected by one-way analysis of variance (ANOVA) (1 factor with ≥3 levels) or two-way ANOVA (≥2 factors with ≥2 levels) followed by pairwise Holm-Šídák post-hoc tests. P ≤ 0.05 was used to determine statistical significance.

## RESULTS

### LC-PUFA supplementation Protects Against TNFα-induced C2C12 Myotube Wasting

To investigate the effects of individual LC-PUFA species with different chain lengths and double bond number/location on myotube atrophy induced by chronic exposure to tumor-derived pro-inflammatory factors, mature C2C12 myotubes were exposed to TNFα with or without individual LC-PUFAs including ARA, EPA, DPA, or DHA for 72 hours. Immunocytochemistry showed that exposure to 100 ng/mL of TNFα induced a significant reduction in myotube diameter (**Figure 1A and B**). Notably, supplementation of the cell culture media with individual LC-PUFAs including ARA, EPA, DPA, and DHA each entirely protected against TNFα-induced myotube atrophy and even resulted in hypertrophy beyond healthy C2C12 myotubes (**Figure 1B**).

**Figure 1.**
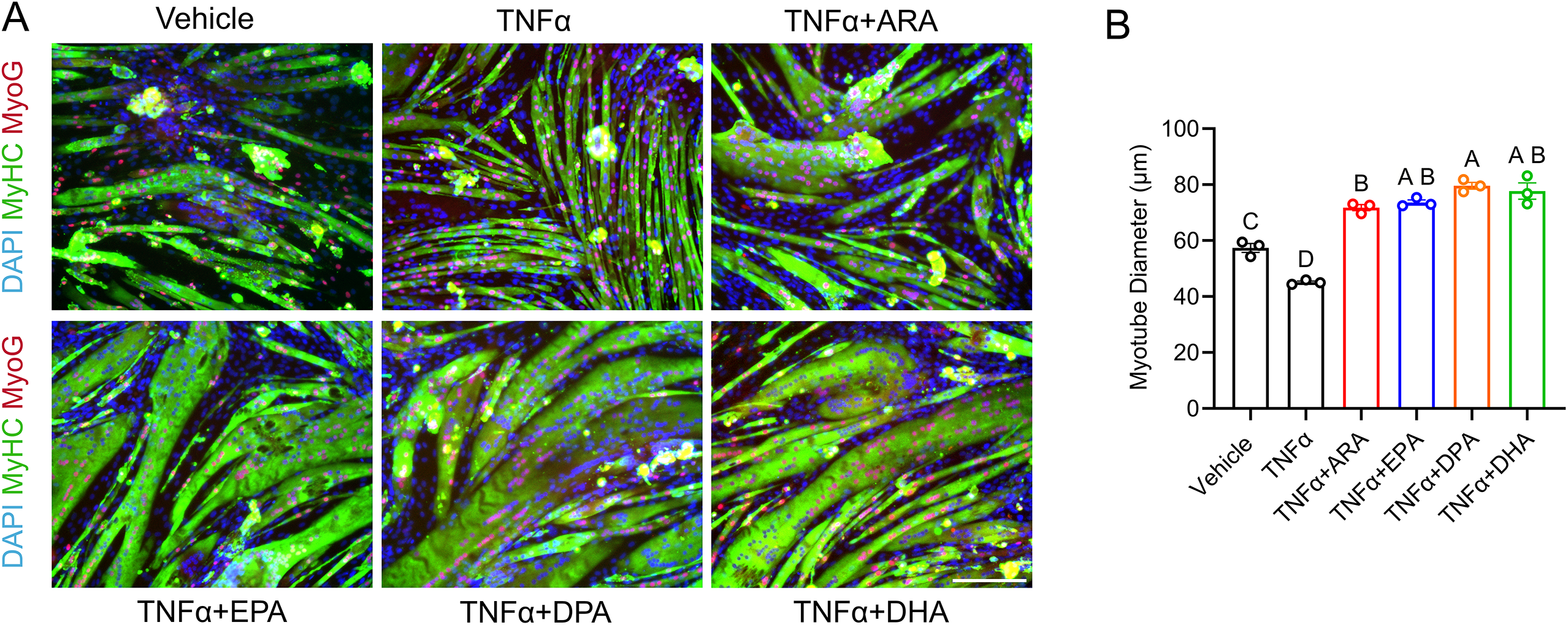
Individual n-3 and n-6 LC-PUFAs prevent TNFα-induced muscle cell wasting. **(A)** Day 3 post- differentiation C2C12 myotubes were treated with 100 ng/mL TNFα in the presence or absence of 25 µM of individual PUFAs including ARA, EPA, DPA, and DHA for 72 h. C2C12 myotubes were fixed and stained for sarcomeric myosin (MF20c, green) and myogenin (F5Dc, red). Nuclei was counter stained by DAPI (blue). (**B)** Quantification of mean myotube diameter in C2C12 myotubes receiving TNFα ± LC-PUFA treatment. Scale bar is 200 μm. Groups labeled with different letters are significantly different from one another, while groups sharing a common letter are not significantly different.

### The Protective Effects of LC-PUFAs against CT26 Cachexia Depend on 15-LOX Enzymatic Activity

Next, we examined the effects of various LC-PUFAs on skeletal muscle atrophy induced by colorectal cancer-cell derived soluble factors. Conditioned cell culture media (CM) samples were obtained from CT26 adenocarcinoma cells cultured *in vitro*. Mature C2C12 myotubes were exposed to CT26 CM in the presence or absence of supplementation with individual n-3 and n-6 LC-PUFAs for 72 h. Treatment of C2C12 myotubes with CT26 CM induced robust muscle cell wasting (**Figure S1**). In contrast, supplementation of CT26 conditioned culture media with individual LC-PUFAs including ARA, EPA, DPA, and DHA, each greatly increased cachectic myotube diameter (**Figure 2A**). We then tested whether the protective effects of LC-PUFAs may depend on the downstream production of bioactive LOX metabolites. C2C12 myotubes exposed to CT26 CM were treated with individual LC-PUFAs in the presence or absence of the pan-LOX inhibitor NDGA or the 15-LOX specific inhibitor BLX-3887. Interestingly, co-treatment with NDGA reversed the protective effects of each LC-PUFAs on CT26 CM cachexia (**Figure 2A and B**). Moreover, when C2C12 myotubes were treated with 15-LOX specific inhibitor BLX-3887, each LC-PUFA tested rather exhibited an overall catabolic effect as evidenced by a significant decrease in myotube diameter below cells treated with CT26 CM alone (**Figure 2A and C**). Gene expression data showed that CT26 CM elevated the expression of pro-inflammatory cytokines including interleukin-6 (IL-6, *Il6)* and monocyte chemoattractant protein 1 (MCP-1, *Ccl2*). Surprisingly however, we observed that ARA, EPA, DPA, and DHA each did not suppress, but rather further increased, CM-induced expression of *Il6* and *Ccl2* (**Figure 2D and E**).

**Figure 2.**
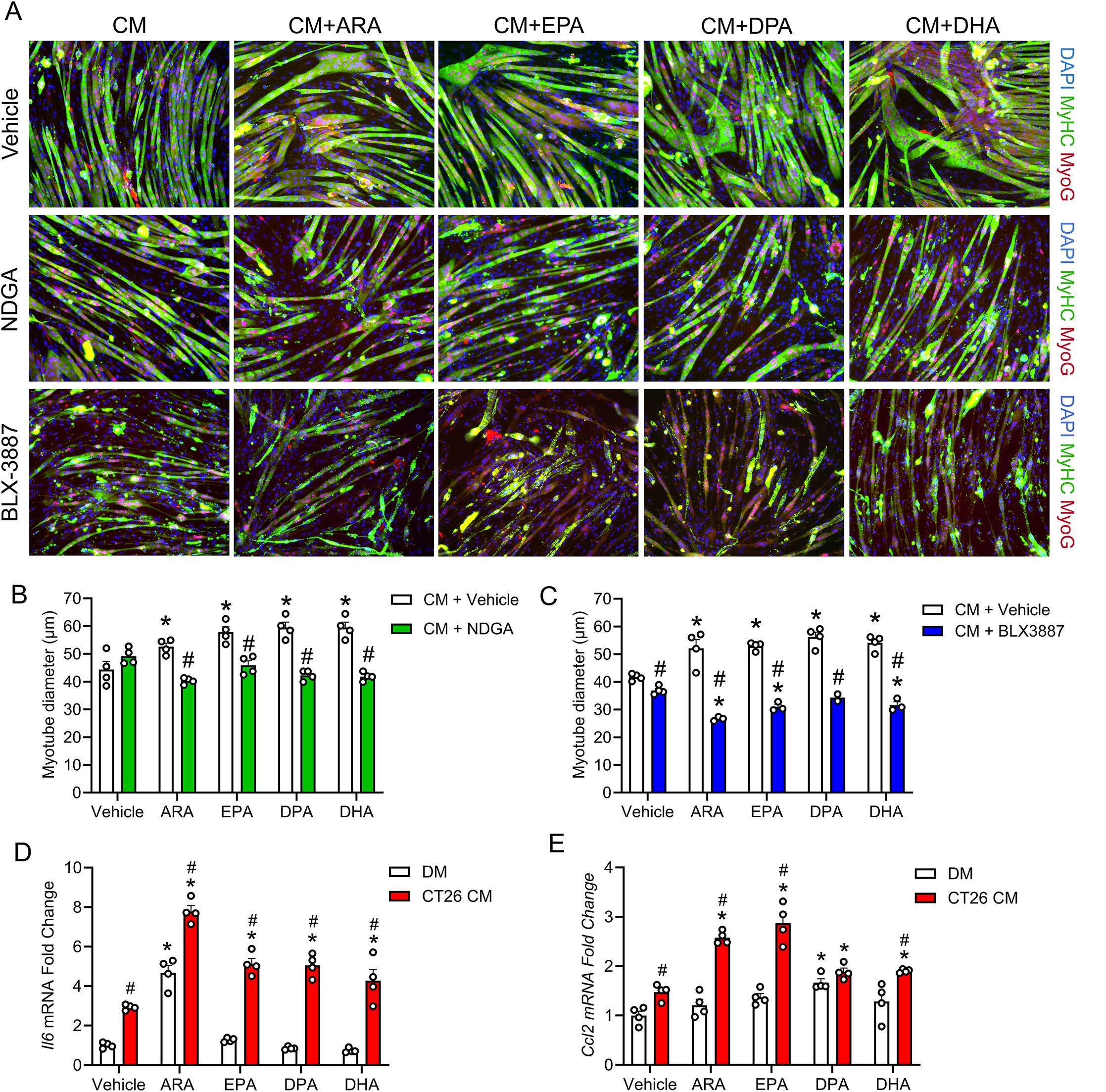
Protective effects of n-3 and n-6 LC-PUFAs against CT26 CM-induced muscle wasting are dependent on 15-LOX activity. Day 3 post-differentiation C2C12 myotubes were treated with CT26 conditioned media (CM) with or without the pan-LOX inhibitor nordihydroguaiaretic acid (NDGA) (50 µM) or the 15-LOX specific inhibitor BLX3887 (10 µM). Then, individual LC-PUFAs including ARA, EPA, DHA, and DPA were spiked into the wells to reach a final concentration of 25 µM. After 72 hours, C2C12 myotubes were fixed in 4% paraformaldehyde (PFA) in preparation for immunocytochemistry analysis. **(A)** C2C12 myotubes were stained for sarcomeric myosin (MF20c, green) and myogenin (F5Dc, red). Nuclei was counter stained by DAPI (blue). **(B-C)** Myotube diameter for C2C12 myotubes with or without NDGA **(B)** or BLX-3887 **(C)** was quantified as described in Methods. Scale bar is 200 μm. *P < 0.05 for difference of PUFA treatments vs. vehicle, and #P < 0.05 for difference between respective LC-PUFA treatments with or without LOX inhibitors. **(D-E)** C2C12 myoblasts were differentiated for 3 days and then exposed to CT26 CM and individual PUFAs including ARA, EPA, DPA, and DHA for 3 hours. RNA was extracted from mature myotubes. mRNA expression fold change of *Il6* **(D)** and *Ccl2* **(E)** in response to CT26 CM and PUFA treatments was shown. *P < 0.05 for effects of PUFA treatments, and #P < 0.05 for difference between DM and CT26 CM.

### Cancer-Muscle Co-Cultures Produce Mature SPMs in Response to LC-PUFA Treatment

To mimic the complex transcellular interactions between tumor and muscle cells in cancer cachexia, an *in vitro* co-culture model was employed to allow the continuous exposure of muscle cells to cancer-derived soluble factors. Co-culture of C2C12 myotubes with CT26 carcinoma cells induced a robust decrease in myotube diameter (>50%) (**Figure 3A and B**). Supplementation with various individual LC-PUFA treatments including ARA, EPA, DPA, and DHA, each significantly increased cachectic myotube diameter, although clearly not to a similar level observed in healthy myotubes (**Figure 3A and B**).

**Figure 3.**
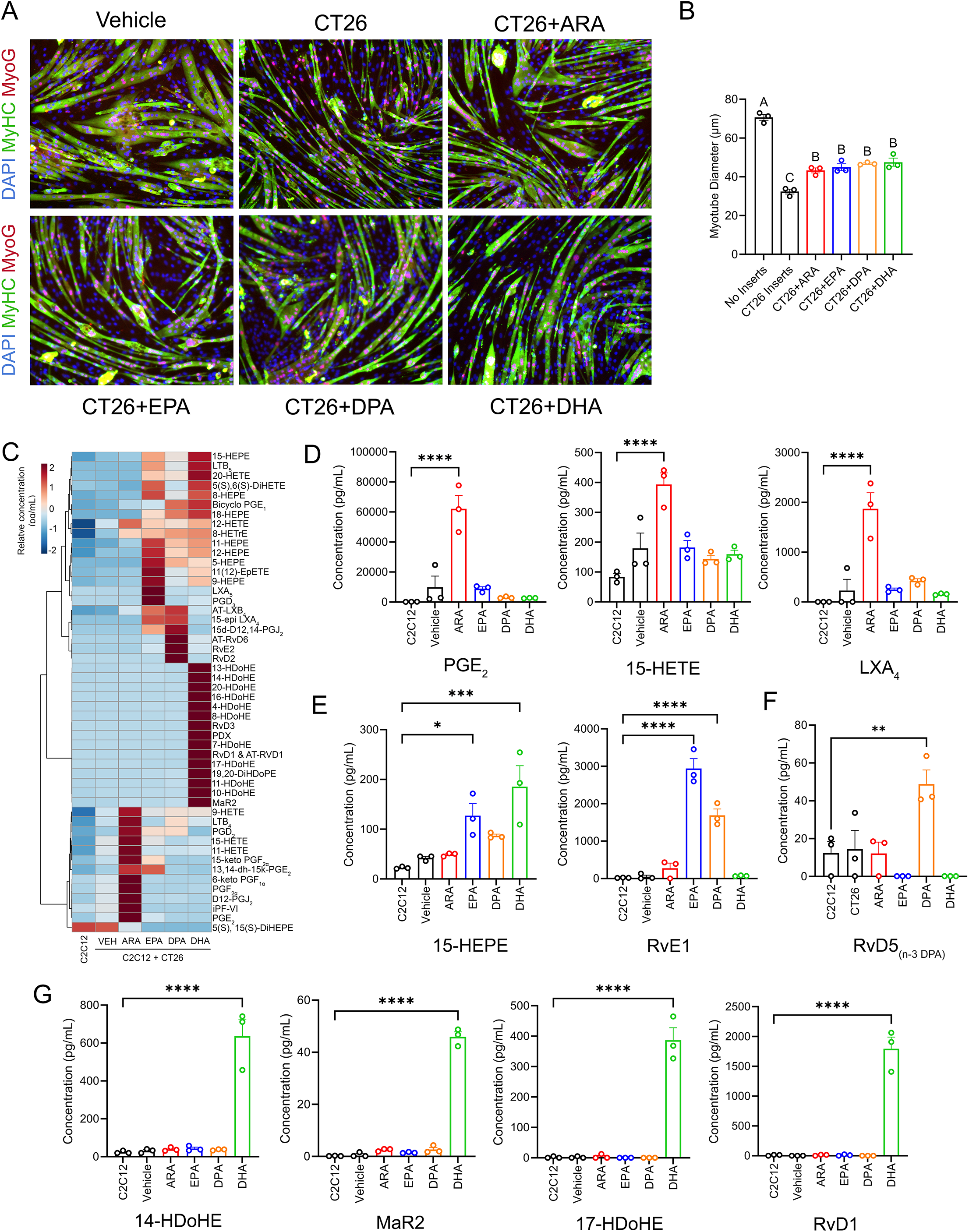
C2C12-CT26 co-cultures produce bioactive lipid metabolites in response to LC-PUFA treatments. Co-cultures of mature C2C12 myotubes and CT26 carcinoma cells were treated with individual LC-PUFAs including ARA, EPA, DPA, and DHA (25 µM). After 72 hours, conditioned media from the lower compartments was collected and C2C12 myotubes were fixed and stained for sarcomeric myosin (MF20c, green) and myogenin (F5Dc, red). Nuclei was counter stained by DAPI (blue) **(A)**. Myotube diameter for C2C12 myotubes in the base plates was quantified as described in Methods **(B)**. Scale bar is 200 μm. Groups labeled with different letters are significantly different from one another, while groups sharing a common letter are not significantly different. **(C- G)** Conditioned media from C2C12-CT26 co-culture was analyzed by targeted Liquid Chromatography-Tandem Mass Spectrometry (LC-MS/MS) to profile lipid metabolites in response to PUFA treatments. **(C)**: A heatmap of the top 50 most differentially regulated lipid mediators detected in conditioned culture media samples from C2C12 myotubes. **(D-G)**: Concentration of representative lipid mediator metabolites downstream of arachidonic acid (ARA) (e.g., PGE_2_, 15-HETE, and LXA^4^) **(D)**, EPA (e.g., 15-HEPE, RvE1) **(E),** DPA (e.g., RvD5_n-3 DPA_) **(F)**, and DHA (e.g., 14-HDoHE, MaR2, 17-HDoHE, and RvD1) **(G)**. Bars show the mean ± SEM of media from 3 wells (biological replicates). P-values were determined by two-tailed unpaired t-tests. ∗p < 0.05, **p<0.01, ***p<0.001, and ****p<0.0001 vs. C2C12 myotubes without PUFAs or CT26 inserts.

To better understand the potential underlying mechanisms responsible for the protective effects of LC-PUFAs in cancer cachexia we employed LC-MS/MS to analyze the metabolipidomic profile of conditioned culture media samples obtained from C2C12-CT26 co-cultures. A complete list of analytes monitored, those which were detected vs. below the limits of detection of the assay, and their average concentration across experimental groups is shown in **Table S1A**. Corresponding signal-to-noise (S/N) ratios for each detected analyte are shown in **Table S1B**. A full list of pairwise comparisons between experimental groups, including fold-change, raw p-values, and Benjamini-Hochberg FDR corrected p-values is shown in **Table S2**. In the absence of LC-PUFA supplementation, the production of 21 lipid mediators was different (raw p<0.05) in C2C12 myotube-CT26 co-cultures compared to healthy C2C12 myotube cultures, including elevated concentrations of leukotriene B_4_ (LTB_4_) and prostaglandin E_2_ (PGE_2_) (**Table S2**). Upon ARA, EPA, DPA, and DHA supplementation, a total of 25, 35, 43, and 49 lipid metabolites were altered (raw p<0.05) in C2C12-CT26 co-cultures, respectively (**Table S2**). In response to LC-PUFA treatments, C2C12-CT26 co-cultures produced markedly higher levels of various downstream bioactive lipid mediators (**Figure 3C**). ARA treatment greatly increased extracellular concentration of downstream ARA metabolites of COX (e.g., PGE_2_), 5-LOX (5-HETE), and 15-LOX (e.g., 15-HETE) (**Figure 3D**). Additionally, we detected markedly elevated levels of the ARA-derived SPM, LXA_4_ in response to ARA supplementation (**Figure 3D**). EPA supplementation induced a higher production of the downstream 15-LOX metabolite of EPA 15-HEPE (**Figure 3E**), the EPA-derived SPMs RvE1 (**Figure 3E**), and RvE2 (**Figure S2D**), and E-series resolvin biosynthetic intermediate 18-HEPE (**Figure S2D**). DHA supplementation resulted in an elevation of primary DHA metabolites of 12-LOX (e.g., 14-HDoHE) and 15-LOX (e.g., 17-HDoHE) (**Figure 3G**). We also detected an increase in mature SPMs derived from DHA including RvD1, RvD3, RvD5, RvD6, MaR2, PD1, PDX, and aspirin-triggered resolvin 3 (AT-RvD3) **(Figure 3G, Figure S2D**). Intriguingly, DHA supplementation also induced an increase in n-3 DPA-derived maresin 1 (MaR1_n-3 DPA_) and protectin D1 (PD1_n-3 DPA_) (**Figure S2C**). Moreover, a higher concentration of 15-HEPE was observed in C2C12-CT26 co-cultures that received DHA supplementation (**Figure 3E**). In response to n-3 DPA treatment, the production of DPA-derived RvD5_n-3DPA_ was increased (**Figure 3F**) as was the EPA-derived SPM RvE1 (**Figure 2E**). We also observed an increased production of DHA-derived RvD2 in response to n-3 DPA supplementation **(Figure S2D**).

### SPMs Differentially Alleviate C2C12 atrophy Induced by TNFα and CT26 Conditioned Media

To determine whether mature SPMs exert protective effects on muscle wasting induced by cancer-associated factors, individual SPMs (100 nM) derived from ARA (LXA_4_), EPA (RvE1), DPA (RvD2_n-3 DPA_), and DHA (RvD1, RvD5, MaR1, and PD1) were administered to C2C12 myotubes exposed to TNFα or CT26 CM for three days. Immunocytochemistry shows that all the mature SPMs tested alleviated skeletal muscle atrophy caused by TNFα (**Figure 4A and B**). However, in the presence of cancer cell conditioned media, only selected D-series SPMs including RvD1, RvD5, and RvD2_n-3 DPA_ significantly increased myotube diameter, indicating a potential differential potency of mature SPMs in mediating cachectic effects induced by distinct tumor-derived factors (**Figure 4C**). We also observed that a 3-hour exposure of TNFα significantly upregulated the expression the pro-inflammatory cytokine interleukin-6 (IL-6, *Il6*) and the major muscle catabolic factor atrogin-1 (*Fbxo32*). Interestingly, while LXA_4_ and PD1 both repressed TNFα-induced *Fbxo32* expression, surprisingly RvD2_n-3DPA_ and RvD5 rather further increased *Fbxo32* expression (**Figure 4E**). Furthermore, RvD1, RvE1, and MaR1 did not significantly impact upon TNFα-induced *Fbxo32* expression. None of the tested SPMs significantly repressed the expression level of *Il6* in response to TNFα stimulation (**Figure 4D**).

**Figure 4.**
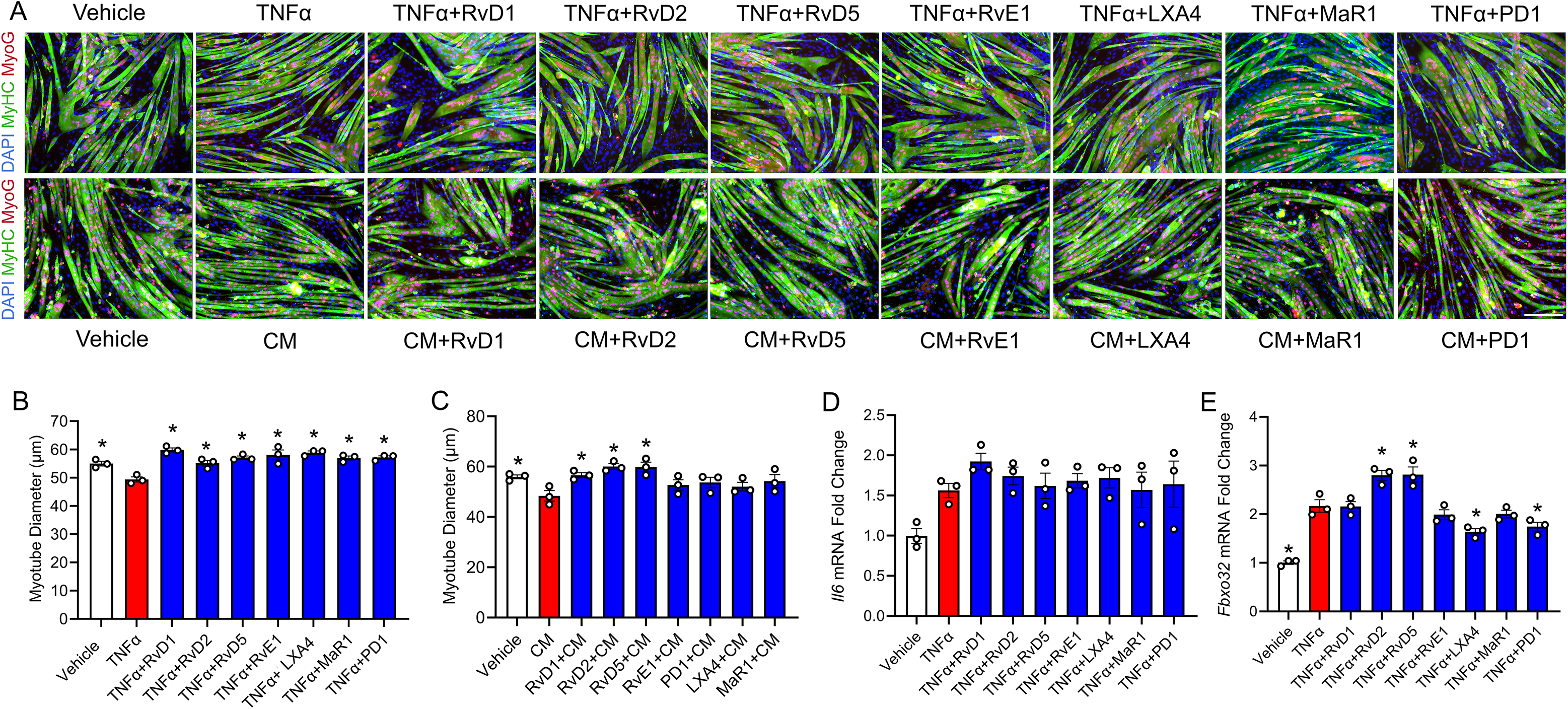
Bioactive SPMs differentially protected against TNFα and CT26 CM-induced muscle wasting in C2C12 cells. **(A)**: Day 3 post-differentiation C2C12 myotubes were treated with 100 ng/mL TNFα (top) or CT26 CM (bottom) in the presence or absence of 100 nM of individual mature SPMs including RvD1, RvD5, MaR1, PD1, RvE1, LXA_4_, and RvD2_n-3 DPA_ for 72 hours. Resulting C2C12 myotubes were fixed and stained for sarcomeric myosin (MF20c, green) and myogenin (F5Dc, red). Nuclei was counter stained by DAPI (blue). **(B- C)** Myotube diameter for C2C12 myotubes receiving TNFα **(B)** or CT26 CM **(C)** was quantified as described in Methods. Scale bar is 200 μm. *P < 0.05 for difference compared to C2C12 myotubes receiving TNFα or CT26 CM alone. **(D-E)** C2C12 myoblasts were differentiated for 3 days and exposed to TNFα (100 ng/mL) with or without individual SPMs including RvD1, RvD5, MaR1, PD1, RvE1, LXA_4_, and RvD2_n-3 DPA_ (100nM) for 3 hours, after which RNA was extracted from mature myotubes. mRNA expression was determined by RT-qPCR and fold change of *Il6* **(D)** and *Fbxo32* **(E)** in response to TNFα exposure and SPM treatments was shown. *P < 0.05 for difference compared to C2C12 myotubes receiving TNFα.

## DISCUSSION

In this study, we examined the potential role of n-3 and n-6 PUFAs and their downstream bioactive SPM metabolites in protecting against skeletal muscle wasting induced by tumor-derived pro-inflammatory factors. Supplementation with various individual LC-PUFAs including n-6 ARA and n-3 EPA, DPA, and DHA each protected cultured myotubes against the atrophic effects of the major tumor-derived pro-inflammatory cytokine TNFα. Similarly, when mature C2C12 myotubes were exposed to conditioned cell culture media derived from CT26 colorectal adenocarcinoma cells, individual PUFAs including ARA, EPA, DPA, and DHA each protected against the atrophic effect of CT26-secreted soluble factors. The remarkable protective effect of each of these LC-PUFAs against tumor cell-derived catabolic factors was found to be dependent on intact 15-LOX activity, inferring a potential causal role of downstream 15-LOX-derived bioactive lipid mediators. Consistently, LC-MS/MS-based metabolipidomic profiling of conditioned media samples obtained from tumor-muscle cell cultures supplemented with LC-PUFAs detected numerous LOX metabolites, including several bioactive LOX-derived SPMs. Finally, treatment of cachectic myotubes with individual synthetic SPMs including ARA-derived LXA_4_, EPA-derived RvE1, DPA-derived RvD2_n-3DPA_, and DHA-derived PD1, RvD1, and RvD5 directly protected skeletal muscle cells against the deleterious effects of exposure TNFα while only select D-series SPMs reversed the catabolic effects of CT26 conditioned media.

The pro-inflammatory cytokine TNFα has been extensively used to model inflammation-driven muscle dysfunction *in vitro* (47). Previous research has demonstrated that n-3 EPA could attenuate TNFα-induced inhibition of myogenic differentiation of C2C12 myoblasts, potentially by upregulating the expression of myogenic differentiation 1 (*Myod1*) and myogenin (*Myog*) (48, 49). Similarly, we observed that n-3 EPA supplementation protected against TNFα induced atrophy of mature C2C12 myotubes. In addition, we show that other longer chain n-3 PUFAs including DPA and DHA also exert a similar protective effect against TNFα- induced muscle wasting. Intriguingly, the major n-6 LC-PUFA ARA also protected cultured myotubes against the catabolic effects of TNFα. ARA is the precursor to PGs and LTs via the action of the COX and 5-LOX pathways, respectively, which play an essential role in skeletal muscle growth and repair (50). Supplementation of ARA has been shown to promote skeletal muscle growth and drive myonuclear accretion (34). Notably however, in the same study, conditioned media from ARA-supplemented C2C12 cells did not appear to exert the same benefits on myotube fusion, indicating possible requirement for continuous ARA exposure and enzymatic metabolism.

We also found that when mature C2C12 myotubes were exposed to conditioned media derived from CT- 26 colorectal carcinoma cells, that individual PUFAs including ARA, EPA, DPA, and DHA each protected against the atrophic effect of CT26-secreted factors. It has been previously shown that colorectal carcinoma cell- secreted soluble factors could induce C2C12 myotube atrophy by increasing IL-6/STAT3 signaling and activating muscle-specific E3 ubiquitin ligases (51, 52). Previous research has also shown that supplementation of EPA-enriched phospholipids could suppress serum TNF-α and IL-6 in tumor-bearing mice (26). However, other studies have shown that supplementation of fish oil containing n-3 PUFAs did not suppress circulatory cytokines despite their ability to attenuate on cachexia (53, 54). In the current study, we found that CT26 CM increased the mRNA expression of pro-inflammatory cytokines including IL-6 (*Il6*) and MCP-1 (*Ccl2*) in C2C12 myotubes. However, perhaps surprisingly, n-3 PUFA supplementation did not suppress the expression *Il6* or *Ccl2*. Rather, EPA, DPA, and DHA each amplified the increase in *Il6* expression in the presence of CT26 CM. Moreover, supplementation of ARA increased the expression of *Il6* both in the presence and absence of CT26 CM. While chronic elevation of IL-6 is commonly associated with cachectic signaling, transiently increased expression of *Il6* in response to PUFA treatments could potentially drive muscle adaptation upon cellular stress (55, 56). Our results suggest a context-based role of IL-6 in PUFA-treated myotubes, which does not necessarily predict muscle wasting morphology.

Interestingly, we found that the anti-cachectic effects of all LC-PUFAs tested was abolished by both pan-LOX inhibitor NDGA and 15-LOX specific inhibitor BLX-3887, indicating the essential role of LOX activity in mediating PUFAs effects. Recent studies have shown that the ablation of leukocyte-type 12/15-LOX in mice models of acute tissue injury resulted in unresolved inflammation and impaired muscle regeneration, demonstrating the indispensable role of 12/15-LOX-derived lipid mediators in myogenesis and muscle repair (41, 57). Moreover, we previously showed that while supplementation with individual LC-PUFAs promoted hypertrophy in wild type primary myotubes, such benefits disappeared in primary myotubes derived from *Alox15^−/−^* mice (41). Moreover, the abolished effects of LC-PUFA-stimulated hypertrophy in *Alox15^−/−^* myotubes was associated with the decreased production of mature SPMs (e.g., RvD2, PDX) (41).

LOX enzymes are a critical enzyme family involved in the biosynthesis of PUFA metabolites including the SPMs, which initiate the resolution phase of muscle inflammation (36). For example, through the enzymatic activity of both 15-LOX and 5-LOX, the n-6 ARA can be converted to LXA_4_ (58). LXA_4_ has been shown to protect against muscle atrophy induced by dexamethasone by attenuating mitochondrial dysfunction (59). The well-known E-series resolvin RvE1 is derived from n-3 EPA through the sequential action of acetylated COX-2 or CYP and 5-LOX (60, 61). RvE1 has been previously shown to repress pro-inflammatory cytokine production and prevent C2C12 myotube atrophy induced by the endotoxin lipopolysaccharide (LPS) (42). Another n-3 LC-PUFA DHA gives rise to the monohydroxy fatty acid intermediate 17-HDoHE through the action of 15-LOX, which is then converted to D-series resolvins (e.g., RvD1) and protectins (e.g., PD1) (62–64). Alternatively, DHA can be catalyzed by the platelet-type12-LOX or the 12-LOX activity of 12/15-LOX to produce 14-HDoHE, which is further converted to maresin (65). We previously showed that RvD1 promotes myofiber regeneration after injury by promoting M2-like phenotype in macrophages and modulating muscle stem cell activity (44). Similarly, PD1 treatment has been shown to promote muscle regeneration in the *mdx* mouse model of Duchenne muscular dystrophy (57). Moreover, exogenous treatment of RvE1 and PD1 promotes neutrophil clearance by macrophage phagocytosis, activating the resolution circuit (66). Despite emerging studies showing the essential role of SPMs in supporting muscle regeneration, to our knowledge no prior published research has examined the influence of SPMs on cancer cachexia. Here, we provide the first evidence that individual LC-PUFAs could protect C2C12 myotubes against CT26-induced atrophy through producing LOX-derived lipid mediators.

In C2C12-CT26 co-cultures, we observed an increased production of many downstream lipid mediators in response to LC-PUFA supplementation. Specifically, ARA supplementation increased the concentration of prostaglandins (e.g., PGE_2_), monohydroxy fatty acids (e.g., 15-HETE), and lipoxins (e.g., LXA_4_). Similarly, EPA increased the level of 18-HEPE and RvE1. DHA induced the production of a wider range of mature SPMs including RvD1, RvD3, AT-RvD3, PD1, PDX, and MaR2. Our data confirms the capacity of C2C12-CT26 co-culture to produce SPMs in response to PUFA substrates, which supports the hypothesis that PUFAs may attenuate CT26-induced muscle wasting specifically through the production of SPMs. Intriguingly, we also observed an increase of RvD2 in response to EPA supplementation, which indicates that EPA might indirectly contribute to DHA-driven metabolites through metabolic interconversion of n-3 PUFAs and potential upregulation of pro-resolving pathway. In addition, there was an increase in 18-HEPE in response to both DPA and DHA supplementation. It has been shown that DHA can potentially be retroconverted to EPA in isolated rat hepatocytes through peroxisomal β-oxidation, and the EPA product can be further elongated to be DPA (67). The supplementation of DHA has been shown to cause an elevated plasma EPA level in humans and rats (68). It is also possible that EPA was released from the membrane upon the cell storage of DHA substrate. However, future research is required to examine the functional capacity of retroconversion from DHA to EPA and/or DPA in muscle cells as well as the change in total fatty acid composition.

Previous research has suggested that CRC is associated with dysregulated lipid metabolism (69). Colorectal tumors from CRC patients produce higher levels of pro-inflammatory lipid mediators (e.g., LTB_4_) while preserving the production of pro-resolving lipid metabolites including 18-HEPE, LXA_4_, and LXB_4_ (69). In the current study, we examined potential shifts in lipid mediator production when C2C12 muscle cells are exposed to cancer-associated factors. In C2C12 myotubes co-cultured with CT26 cells, the production of several lipid metabolites (e.g., LTB_4_, PGE_2_) was elevated compared to healthy C2C12 myotubes.

We found that direct treatment of a range of mature SPMs fully protected C2C12 myotubes against TNFα- induced atrophy. However, when exposed to CT26 CM, mature SPMs exerted differential potential in rescuing the cachectic effect. CT26 CM could possess a complex tumor secretome including cytokines (e.g., IL-6, TNFα, IL-1β), chemokines (e.g., CXCL1), and extracellular vesicles (e.g., exosomes) (70–72). The integrated factors could contribute to a systemic atrophy beyond the rescuing capacity of a single SPM treatment. Moreover, we found that single dose of SPMs did not downregulate the mRNA expression of *Il6* induced by TNFα, and only LXA_4_ and PD1 significantly repressed the mRNA expression of Atrogin-1 (*Fbxo32*), which is a muscle-specific E3 ubiquitin ligase often involved in cancer-induced muscle wasting (73). Our results suggests that SPMs could attenuate TNFα-induced muscle wasting through mechanisms other than modulating cytokine signaling or protein degradation, and the preserved muscle morphology might not necessarily require changes in cachexia-associated markers. Although previous research shows that RvD1 could suppress *Il6* expression in C2C12 exposed to LPS, how SPMs mediate cytokine expression and proteolysis pathway in cancer cachexia remains unclear (44). Further research could look into the role of SPMs in anabolic signaling, immune environment, and muscle repair in CRC cachexia.

One limitation of our study is the current lack of *in vivo* model to resemble the complex systemic environment of CRC cachexia and investigate how LC-PUFA supplementation regulates immune-muscle interactions. As such, our future research will examine the role of dietary supplementation of n-3 and n-6 LC-PUFAs in mediating CRC cachexia in *in vivo* animal models. Moreover, future studies could examine other potential pathways involved in CRC cachexia, such as protein synthesis/degradation pathway, muscle energy metabolism, and mitochondrial function. Finally, future research could explore whether the effects of the frequency, dose, and duration of administration of LC-PUFAs and SPMs in models of cancer cachexia.

## CONCLUSION

In the current study, we investigated the effect of supplementation with individual n-3 and n-6 LC-PUFAs in CRC-induced skeletal muscle wasting *in vitro*. We found that LC-PUFAs including ARA, EPA, DPA, and DHA each individually alleviated C2C12 myotube atrophy induced by CT26 conditioned media, and the attenuating effects of LC-PUFAs was dependent on the enzymatic activity of 15-LOX. We show that tumor-muscle cell co-cultures exposed to individual PUFA treatments produce a wide range of downstream lipid mediator including mature SPMs. All SPMs tested protected C2C12 myotubes against the deleterious effects of exposure to TNFα. However, direct treatments of mature SPMs in C2C12 myotubes exposed to CT26 CM exerted differential protective effects, with only D-series SPMs including RvD1, RvD5, and RvD2_n-3 DPA_ preventing muscle atrophy.

## AUTHOR CONTRIBUTIONS

**Xinyue Lu:** Conceptualization, Methodology, Validation, Formal analysis, Investigation, Writing – original draft, Writing – review & editing, Visualization. **Krishna Rao Maddipati:** Methodology, Investigation, Resources, Writing – review & editing. **James F. Markworth:** Conceptualization, Methodology, Software, Formal analysis, Resources, Writing – review & editing, Visualization, Supervision, Project administration, Funding acquisition.

## ACKNOWLEDGEMENT

The MF20 (developed by Fischman, D.A.) and F5D (developed by Wright, W. E.) monoclonal antibodies were obtained from the Developmental Studies Hybridoma Bank (DSHB), created by the NICHD of the NIH and maintained at The University of Iowa, Department of Biology, Iowa City, IA 52242.

---

**Figure S1.**
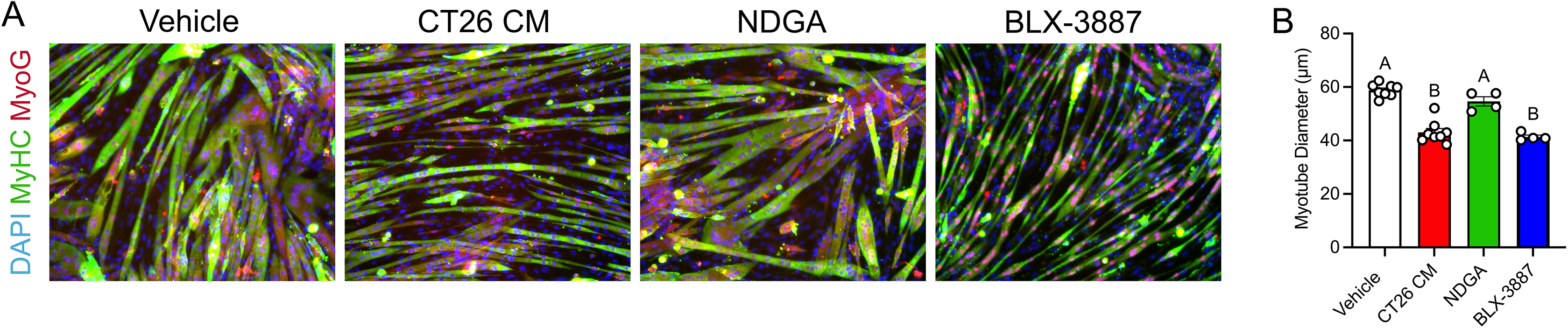
CT26 CM induces muscle wasting in C2C12 cells. **(A)**: Day 3 post-differentiation C2C12 myotubes were treated with fresh DM, CT26 CM, DM containing 50 µM NDGA, or 10 µM BLX-3887 for 72 hours. C2C12 myotubes were fixed and stained for sarcomeric myosin (MF20c, green) and myogenin (F5Dc, red). Nuclei was counter stained by DAPI (blue). **(B)**: Myotube diameter for C2C12 myotubes was quantified as described in Methods. Groups labeled with different letters are significantly different from one another, while groups sharing a common letter are not significantly different.

**Figure S2.**
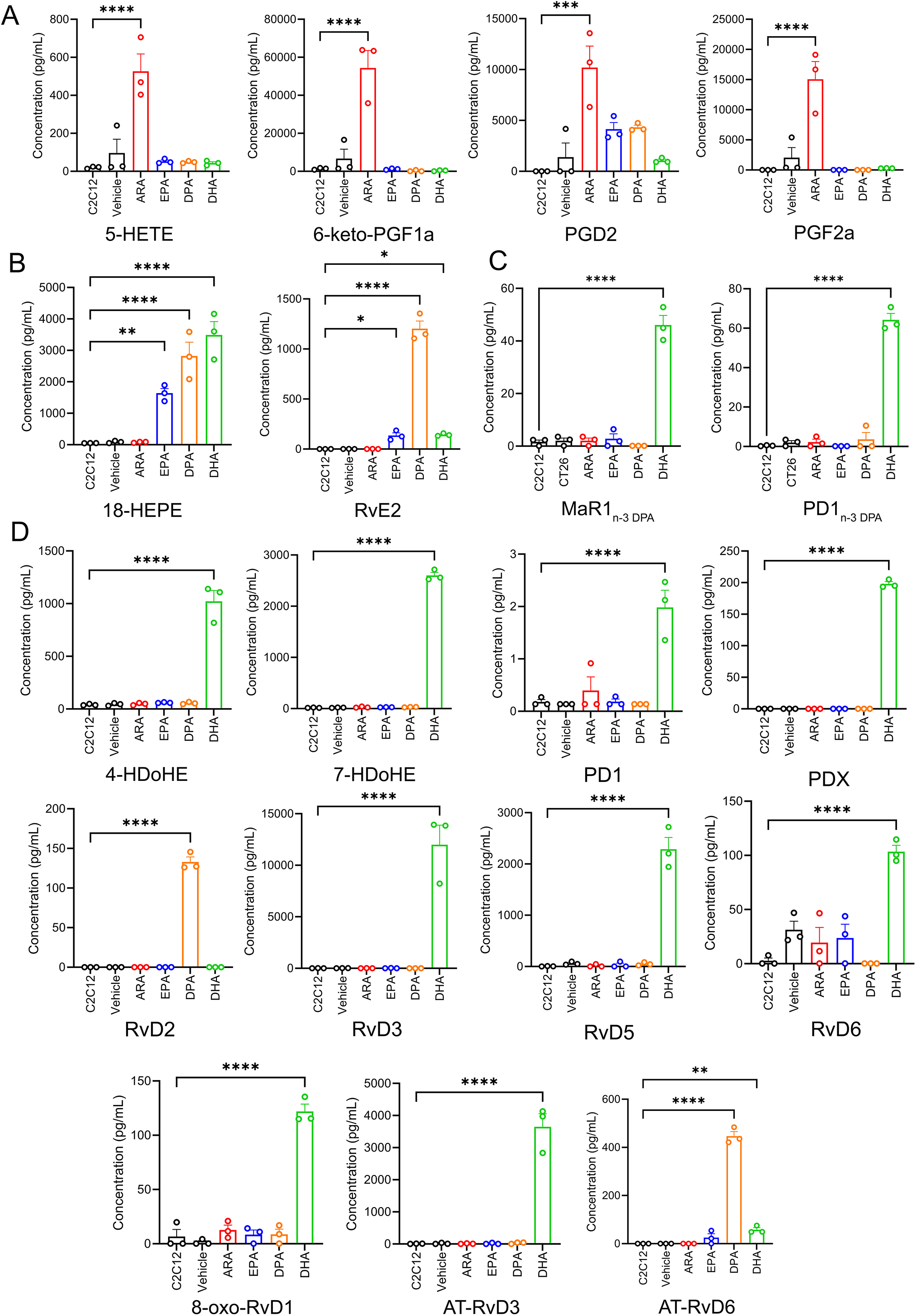
Representative bioactive lipid metabolites produced by C2C12-CT26 co-culture in response to LC-PUFA treatments. Concentration of lipid mediator metabolites downstream of arachidonic acid (ARA) (e.g., 5-HETE, 6-keto-PGF_1α_, PGD_2_, and PGF_2α_) **(A)**, EPA (e.g., 18-HEPE, RvE2) **(B),** DPA (e.g., MaR1_n-3 DPA_, PD1_n- 3 DPA_) **(C)**, and DHA (e.g., 4-HDoHE, 7-HDoHE, PD1, PDX, RvD2, RvD3, RvD5, RvD6, AT-RvD3, AT-RvD6) **(D)** Bars show the mean ± SEM of media from 3 wells (biological replicates). P-values were determined by two-tailed unpaired t-tests. ∗p < 0.05, **p<0.01, ***p<0.001, and ****p<0.0001 vs. C2C12 myotubes without PUFAs or CT26 inserts.

